# Physiological determinants of the increase in oxygen consumption during exercise in individuals with stroke

**DOI:** 10.1101/641522

**Authors:** Kazuaki Oyake, Yasuto Baba, Nao Ito, Yuki Suda, Jun Murayama, Ayumi Mochida, Kunitsugu Kondo, Yohei Otaka, Kimito Momose

## Abstract

**Background:** Understanding the physiological limitations of the increase in oxygen consumption (V̇O_2_) during exercise is essential to improve cardiorespiratory fitness in individuals with stroke. However, the physiological determinants of the increase in V̇O_2_ during exercise have not been examined using multivariate analysis in individuals with stroke. This study aimed to identify the physiological determinants of the increase in V̇O_2_ during a graded exercise in terms of the respiratory function, cardiac function, and ability of skeletal muscles to extract oxygen.

**Methods:** Eighteen individuals with stroke (60.1 ± 9.4 years of age, 67.1 ± 30.8 days poststroke) underwent a graded exercise test for the assessment of cardiorespiratory response to exercise. The increase in V̇O_2_ from rest to ventilatory threshold and that from rest to peak exercise were measured as a dependent variable. The increases in respiratory rate, tidal volume, heart rate, stroke volume, and arterial-venous oxygen difference from rest to ventilatory threshold and those from rest to peak exercise were measured as independent variables.

**Results:** From rest to ventilatory threshold, the increases in heart rate (β = 0.546) and arterial-venous oxygen difference (β = 0.398) were significant determinants of the increase in V̇O_2_ (adjusted R^2^ = 0.703, p < 0.001). From rest to peak exercise, the increases in tidal volume (β = 0.611) and heart rate (β = 0.353) were significant determinants of the increase in V̇O_2_ (adjusted R^2^ = 0.702, p < 0.001).

**Conclusion:** V̇O_2_ is well-known to increase nearly linearly with increasing heart rate; however, our results suggest that arterial-venous oxygen difference and tidal volume are also significant physiological determinants of the increase in V̇O_2_ from rest to ventilatory threshold and that from rest to peak exercise, respectively. Our findings could potentially contribute to the development of appropriate therapies to improve cardiorespiratory fitness in individuals with stroke.

## Introduction

Individuals with stroke have reduced cardiorespiratory fitness compared with age- and sex-matched healthy individuals [1, 2]. Cardiorespiratory fitness reduction is potentially related to walking disability [3, 4], poor cognitive performance [5], and limitations in activities of daily living in individuals with stroke [6-8]. Low levels of cardiorespiratory fitness following stroke may lead to avoidance of physical activity, which causes further deconditioning [9, 10]. Therefore, understanding the physiological limitations of cardiorespiratory fitness in individuals with stroke is essential for the development of appropriate therapies to improve physical activity levels and to prevent further deconditioning.

Oxygen consumption (V̇O_2_) at ventilatory threshold and that at peak exercise measured during a graded exercise test are used to assess cardiorespiratory fitness in individuals with stroke [1, 11-13]. The physiological factors that potentially limit V̇O_2_ at peak exercise are respiratory and cardiac functions to supply oxygen and the ability of skeletal muscles to extract oxygen [12, 14, 15]. In healthy adults, V̇O_2_ at peak exercise seems to be limited primarily by cardiac function [14, 15]. Previous studies [12, 16-18] reported that tidal volume, heart rate, and arterial-venous oxygen difference at peak exercise are significantly lower in individuals with stroke than those in age- and sex-matched healthy adults, which may lead to the deterioration of cardiorespiratory fitness after stroke. Tomczak et al. [18] reported a significant difference between individuals with stroke and healthy adults in V̇O_2_ at peak exercise, but not in V̇O_2_ at rest. Thus, identifying the determinants of the increase in V̇O_2_ during exercise contributes to understanding the physiological limitations of cardiorespiratory fitness in individuals with stroke. However, the physiological determinants of the increase in V̇O_2_ during exercise have not been examined using multivariate analysis in individuals with stroke.

Cross-sectional and longitudinal studies found the relationship between V̇O_2_ and arterial-venous oxygen difference at peak exercise in individuals with stroke [17, 19]. Therefore, we hypothesized that arterial-venous oxygen difference is a primary physiological determinant of the increase in V̇O_2_ during exercise in these individuals. In this study, we aimed to explore the physiological determinants of the increase in V̇O_2_ from rest to ventilatory threshold and that from rest to peak exercise in individuals with stroke.

## Methods

### Study design

This study used a cross-sectional observational design. The study protocol was approved by the appropriate ethics committees of the Tokyo Bay Rehabilitation Hospital (approval number: 172-2) and the Shinshu University (approval number: 3813). All participants provided written informed consent prior to study enrollment. The study was conducted according to the Declaration of Helsinki of 1964, as revised in 2013.

### Participants

Participants were recruited from a convalescent rehabilitation hospital between November 2017 and November 2018. The inclusion criteria for the study were as follows: (1) age 40–80 years, (2) being within 180 days after first-ever stroke, (3) ability to maintain a target cadence of 50 rpm during exercise, and (4) a Mini-Mental State Examination score [20] of 24 or more. The exclusion criteria were as follows: (1) limited range of motion and/or pain that could affect the exercise test, (2) unstable medical conditions such as unstable angina, uncontrolled hypertension, and tachycardia, (3) use of beta-blocker, and (4) any comorbid neurological disorder.

### Exercise testing

Participants were instructed to refrain from eating for 3 hours and to avoid caffeine and vigorous physical activity for at least 6 and 24 hours, respectively, before the exercise test [21]. All participants performed a symptom-limited graded exercise test on a recumbent cycle ergometer (Strength Ergo 240; Mitsubishi Electric Engineering Co., Ltd., Tokyo, Japan) that can be precisely load-controlled (coefficient of variation, 5%) over a wide range of pedaling resistance (0-400 W). The distance from the seat edge to pedal axis was adjusted so that the participant’s knee flexion angle was 20° when extended maximally. The backrest was set at 20° reclined from the vertical position. Additional strapping was attached to secure the paretic foot to the pedal as needed. Following a 3-min of rest period (in sitting position) on the recumbent cycle ergometer to establish a steady state, a warm-up was performed at 0 W for 3 min followed by 10 W increments every minute [21, 22]. Participants were instructed to maintain a target cadence of 50 rpm throughout the exercise [21, 22]. Blood pressure was monitored every minute from the non-paretic arm using an automated system (Tango; Sun Tech Medical Inc., NC, USA). The test was terminated if the participants showed signs of angina, dyspnea, inability to maintain cycling cadence more than 40 rpm, hypertension (more than 250 mmHg systolic or more than 115 mmHg diastolic), or drop in systolic blood pressure more than 10 mmHg despite an increase in work load [22, 23]. Participants provided their ratings of perceived exertion (6 = no exertion at all, 20 = maximal exertion) [24] for dyspnea and leg effort at the end of the test. Work rate at peak exercise was defined as the peak wattage on test termination [19]. To identify whether maximal effort was reached during the exercise test, at least 1 of the following criteria had to be met: (1) V̇O_2_ increased less than 150 mL·min^−1^ for more than 1 min despite increased work rate, (2) respiratory exchange ratio achieved greater than 1.10, 1. (3) or heart rate achieved 85% of the age-predicted maximal heart rate (220 minus age) [23, 25].

Physiological variables were measured at rest and continuously during exercise test. V̇O_2_, respiratory rate, and tidal volume were measured on a breath-by-breath basis using an expired gas analyzer (Aerosonic AT-1100; ANIMA Corp., Tokyo, Japan). Carbon dioxide output, the ventilatory equivalents of oxygen and carbon dioxide, and the end-tidal oxygen and carbon dioxide fractions were also measured using the expired gas analyzer to determine the ventilatory threshold. Heart rate and stroke volume were measured on a beat-by-beat basis using a noninvasive impedance cardiography device (Task Force Monitor model 3040i; CN Systems Medizintechnik GmbH., Graz, Austria) [26-28]. For calculating arterial-venous oxygen difference, measurement values of physiological variables were interpolated to 1-s intervals, time-aligned, and averaged into 5-s bins [18]. Arterial-venous oxygen difference was calculated as the ratio between V̇O_2_ and the product of heart rate and stroke volume.

Physiological variables at rest were defined as the average value obtained during 1 min before exercise onset, and those at peak exercise were defined as the average value obtained during the last 30 s of exercise test [18, 21]. The ventilatory threshold was determined using a combination of the following criteria: (1) the point where the ventilatory equivalent of oxygen reaches its minimum or starts to increase, without an increase in the ventilatory equivalent of carbon dioxide; (2) the point at which the end-tidal oxygen fraction reaches a minimum or starts to increase, without a decline in the end-tidal carbon dioxide fraction; and (3) the point of deflection of carbon dioxide output versus V̇O_2_ [29]. The first two methods were prioritized in case the three methods presented different results [30, 31]. The ventilatory threshold was determined as an average based on the values provided by two independent raters (NI and YS), when the difference in the V̇O_2_ values of the corresponding points as determined by the two raters was less than 100 mL·min^−1^ [31, 32]. In case of any discrepancy, a third experienced rater (KO) judged the point, and the ventilatory threshold was taken as the average of the two closest values [30, 31].

### Statistical analysis

The G Power computer program version 3.1.9.2 (Heinrich Heine University, Dusseldorf, Germany) [33] was used to calculate the sample size required for multiple regression analysis. For multiple regression analyses, if up to five variables (respiratory rate, tidal volume, heart rate, stroke volume, and arterial-venous oxygen difference) are modeled at an effect size of 0.49 (very large) at an α level of 0.05 and power of 0.80, a minimum of 12 participants are required [33, 34].

Normality of distribution was tested using the Shapiro-Wilk test. One-way repeated-measures analysis of variance or Friedman test with exercise period as a factor was used to examine whether physiological variables change during exercise. Post hoc analyses were performed using the Bonferroni multiple comparison test.

The increase in V̇O_2_ from rest to ventilatory threshold and that from rest to peak exercise were calculated as the dependent variables. Pearson’s product moment correlation coefficient or Spearman’s rank correlation coefficient was used to test the correlations between the increases from rest to ventilatory threshold in V̇O_2_ and other physiological variables and between the increases from rest to peak exercise in V̇O_2_ and other physiological variables. Those variables that were significantly correlated with the increase in V̇O_2_ during exercise testing were then entered into the stepwise multiple regression analysis to determine the physiological limitations of the increase in V̇O_2_ with considering multicollinearity. Statistical analyses were performed using the Statistical Package for the Social Sciences software version 24.0 (International Business Machines Corp., NY, USA). Any p values less than 0.05 were considered statistically significant.

## Results

A flow diagram of study participants is shown in Fig 1. Eighteen individuals with stroke participated in the study. Table 1 shows the characteristics of the participants.

**Fig 1.**
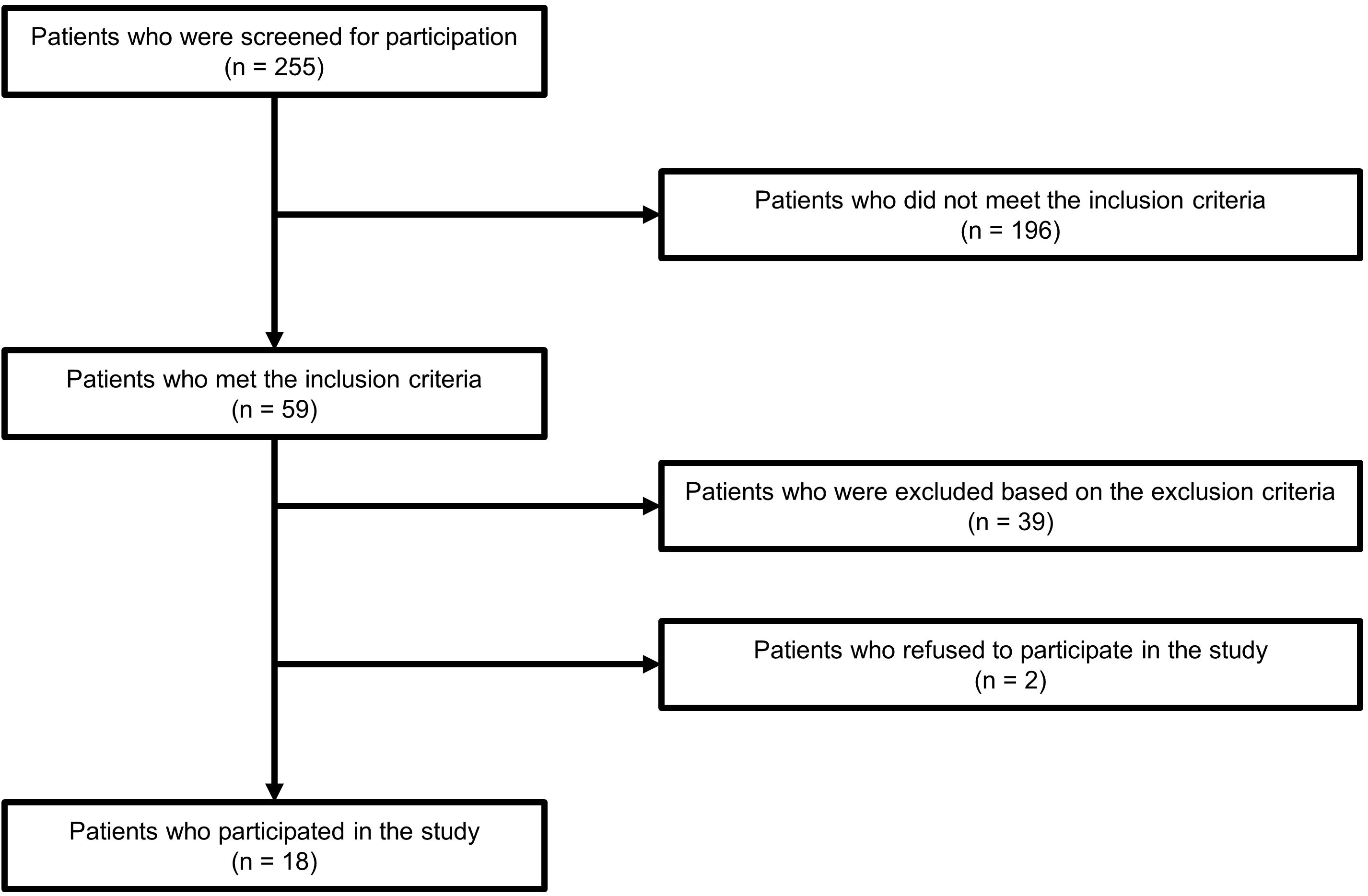
Flow diagram of study participants.

**Table 1.**
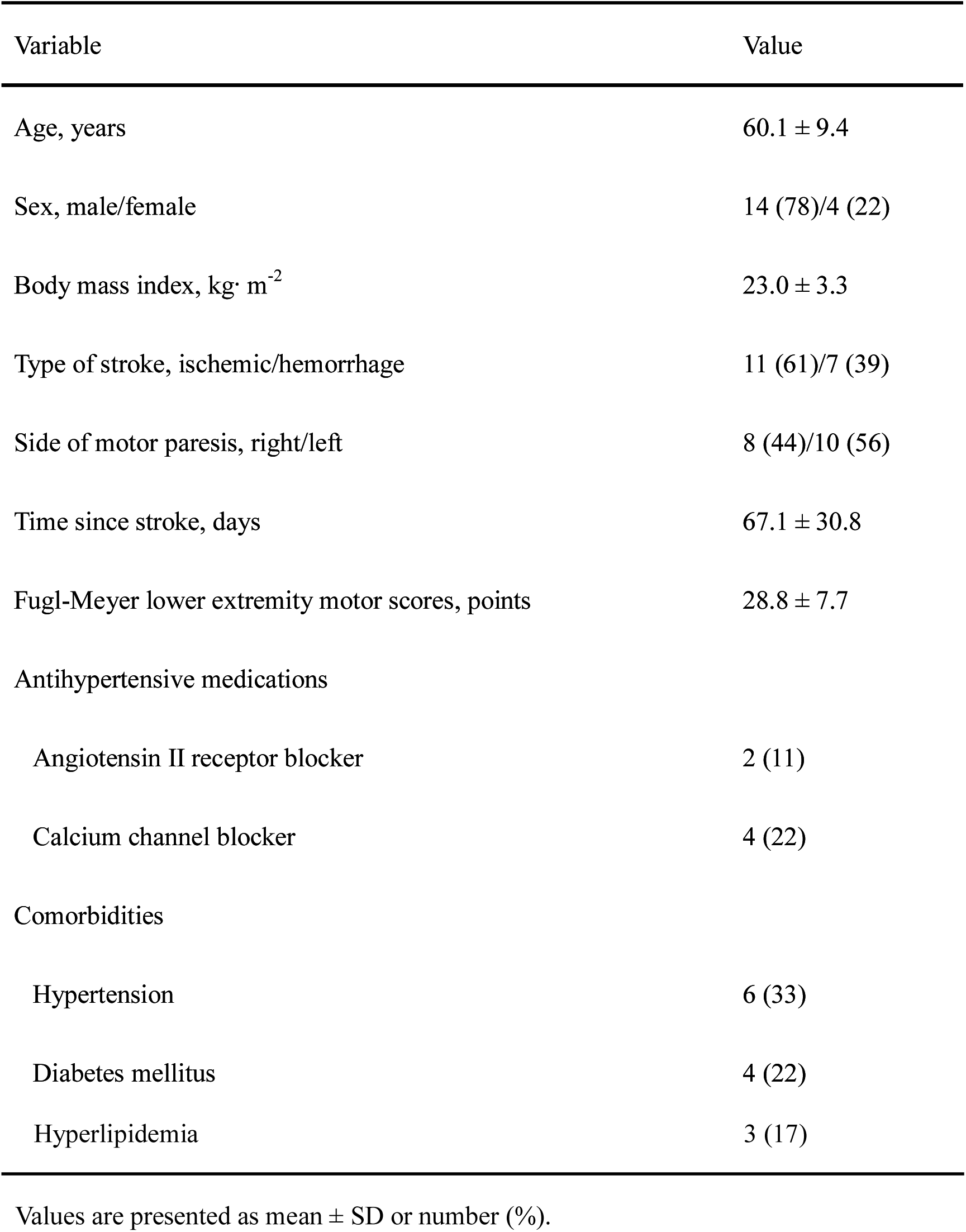
Characteristics of participants

No significant adverse events occurred during or after exercise test. All participants stopped their exercise test due to inability to maintain cycling cadence more than 40 rpm. In addition, all participants met at least one of the criteria for reaching maximal effort. Median (interquartile range) values of the ratings of perceived exertion for dyspnea and leg effort at the end of the test were 13 (13–15) and 15 (13–15), respectively. Mean ± standard deviation of work rate at peak exercise was 69.4 ± 30.6 W.

Measurement values at rest, ventilatory threshold, and peak exercise are shown in Table 2. We observed a main effect of exercise period on all physiological variables (p < 0.001). All physiological variables at ventilatory threshold were significantly higher than those at rest (p < 0.001). From ventilatory threshold to peak exercise, V̇O_2_, respiratory rate, tidal volume, and heart rate significantly increased (mean difference = 198.6, 95% confidence interval [CI] = 95.4, 301.7, and p < 0.001; mean difference = 7.5, 95% CI = 3.2, 11.7, and p < 0.001; mean difference = 0.21, 95% CI = 0.08, 0.33, and p = 0.002; mean difference = 18.7, 95% CI = 10.2, 21.3, and p < 0.001, respectively), whereas no significant changes in stroke volume and arterial-venous oxygen difference were found (mean difference = −1.9, 95% CI = −6.9, 3.0, and p = 0.945; mean difference = 0.74, 95% CI = −0.22, 1.70, and p = 0.172, respectively).

**Table 2.** Physiological variables at rest, ventilatory threshold, and peak exercise

| Variable | Rest | Ventilatory threshold | Peak exercise | p value |
| --- | --- | --- | --- | --- |
| $\dot{V}O_2$, mL·min<sup>-1</sup> | 253.1 ± 57.7 | 899.7 ± 224.8<sup>*</sup> | 1098.3 ± 330.3<sup>†, ‡</sup> | < 0.001 |
| Respiratory rate, breaths·min<sup>-1</sup> | 18.5 ± 2.8 | 25.2 ± 4.5<sup>*</sup> | 32.6 ± 8.5<sup>†, ‡</sup> | < 0.001 |
| Tidal volume, L | 0.46 ± 0.11 | 1.07 ± 0.23<sup>*</sup> | 1.28 ± 0.30<sup>†, ‡</sup> | < 0.001 |
| Heart rate, bpm | 76.0 ± 9.2 | 109.9 ± 13.6<sup>*</sup> | 128.6 ± 17.5<sup>†, ‡</sup> | < 0.001 |
| Stroke volume, mL | 55.0 ± 7.6 | 73.0 ± 10.4<sup>*</sup> | 71.0 ± 9.4<sup>†</sup> | < 0.001 |
| Arterial-venous oxygen difference, mL·100 mL<sup>-1</sup> | 6.2 ± 1.6 | 11.2 ± 1.8<sup>*</sup> | 12.0 ± 2.4<sup>†</sup> | < 0.001 |

Values are presented as mean ± SD.
A p value represents a significant main effect of exercise period.
<sup>\*</sup>, a significant difference between ventilatory threshold and rest (p < 0.001).
<sup>†</sup>, a significant difference between peak exercise and rest (p < 0.001).
‡, a significant difference between peak exercise and ventilatory threshold ( $p < 0.001$ ).

From rest to ventilatory threshold, correlations between the increases in V̇O_2_ and other physiological variables are shown in Fig 2. The increase in V̇O_2_ was significantly correlated with the increases in tidal volume (r = 0.620; 95% CI = 0.215, 0.843; and p = 0.006) (Fig 2b), heart rate (r = 0.804; 95% CI = 0.540, 0.924; and p < 0.001) (Fig 2c), and arterial-venous oxygen difference (r = 0.752; 95% CI = 0.440, 0.902; and p < 0.001) (Fig 2e). Stepwise multiple regression analysis revealed that the increases in heart rate (β = 0.546) and arterial-venous oxygen difference (β = 0.398) were the significant determinants for the increase in V̇O_2_ (adjusted R^2^ = 0.703, p < 0.001) (Table 3).

**Fig 2.**
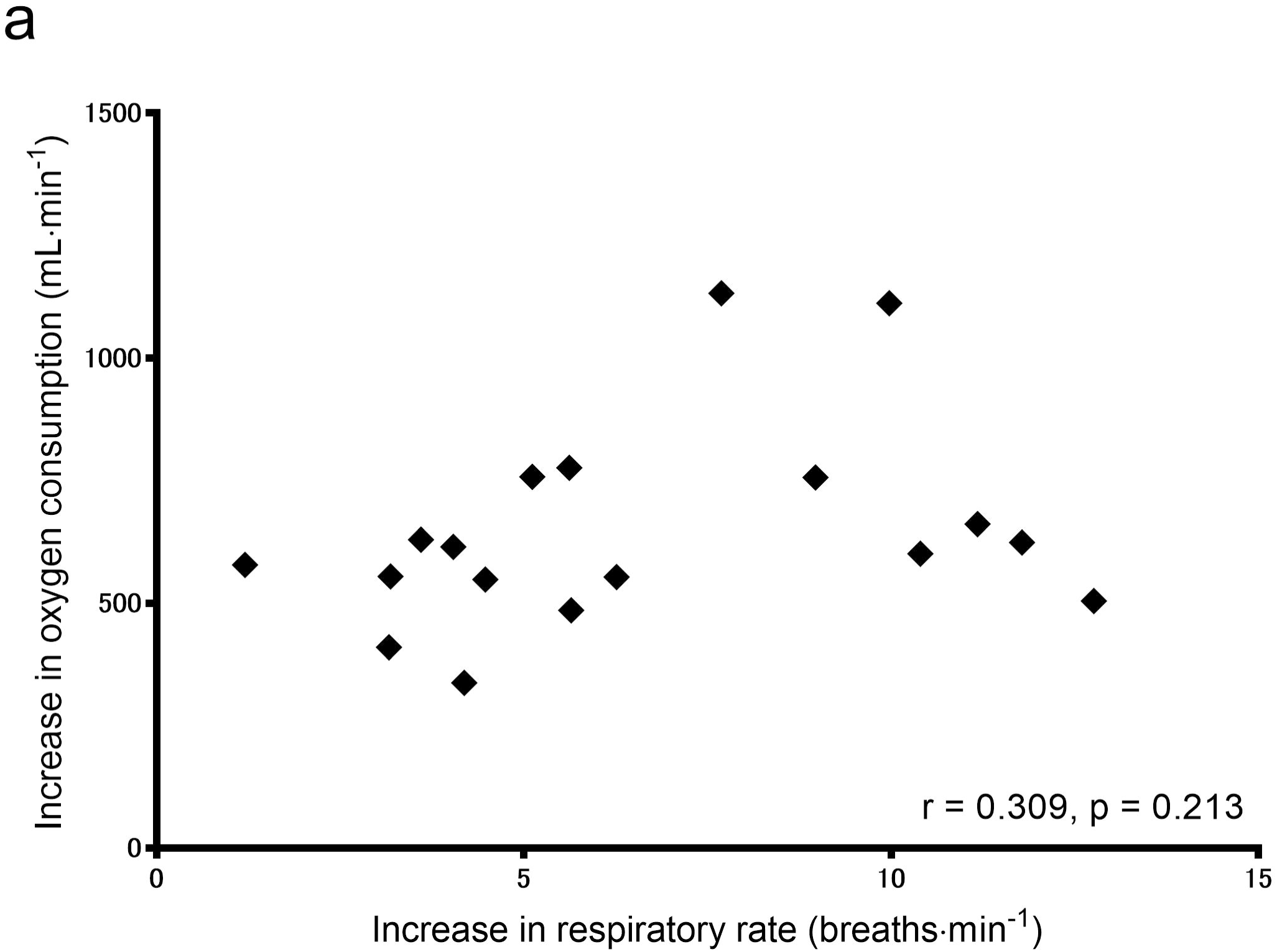
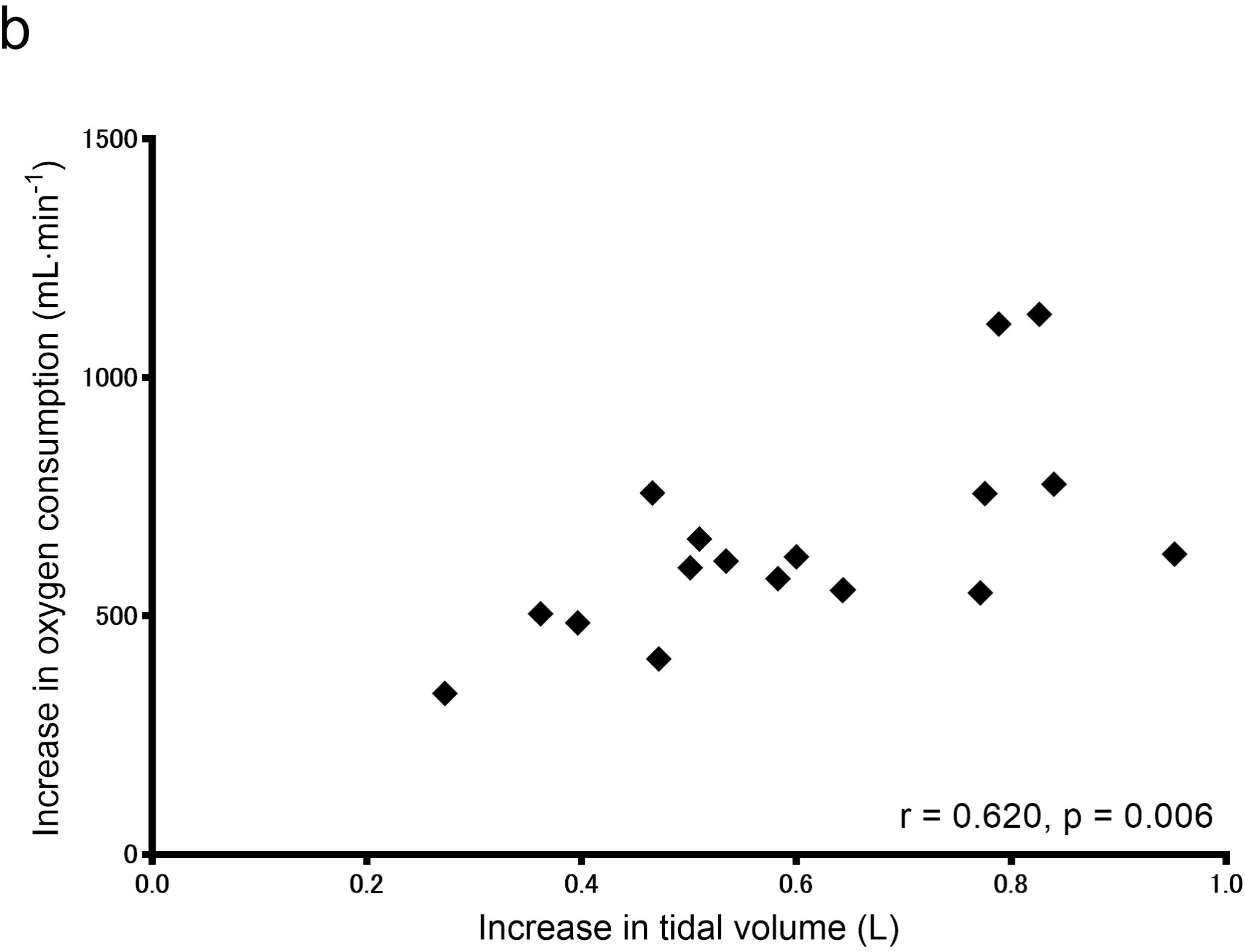
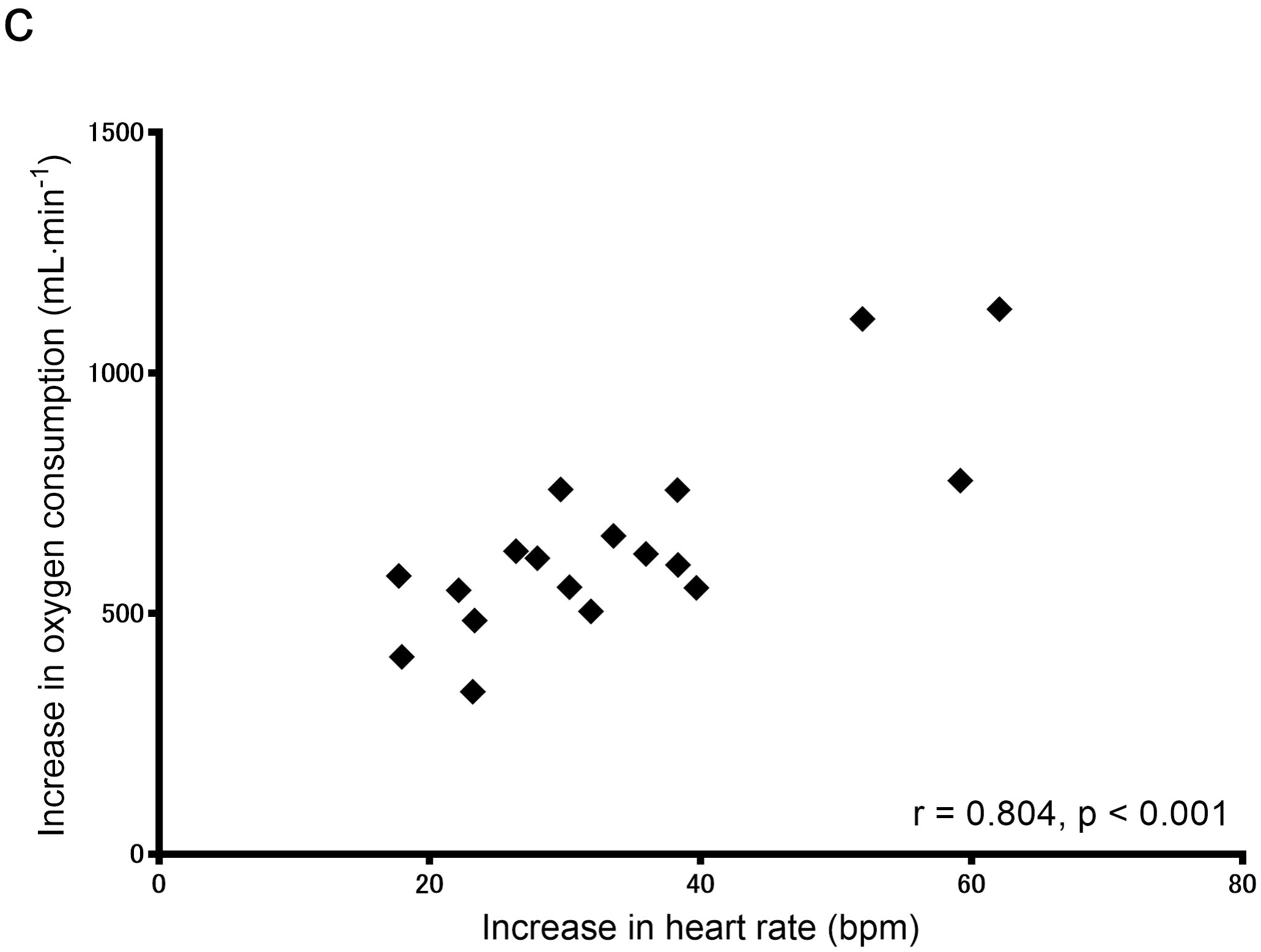
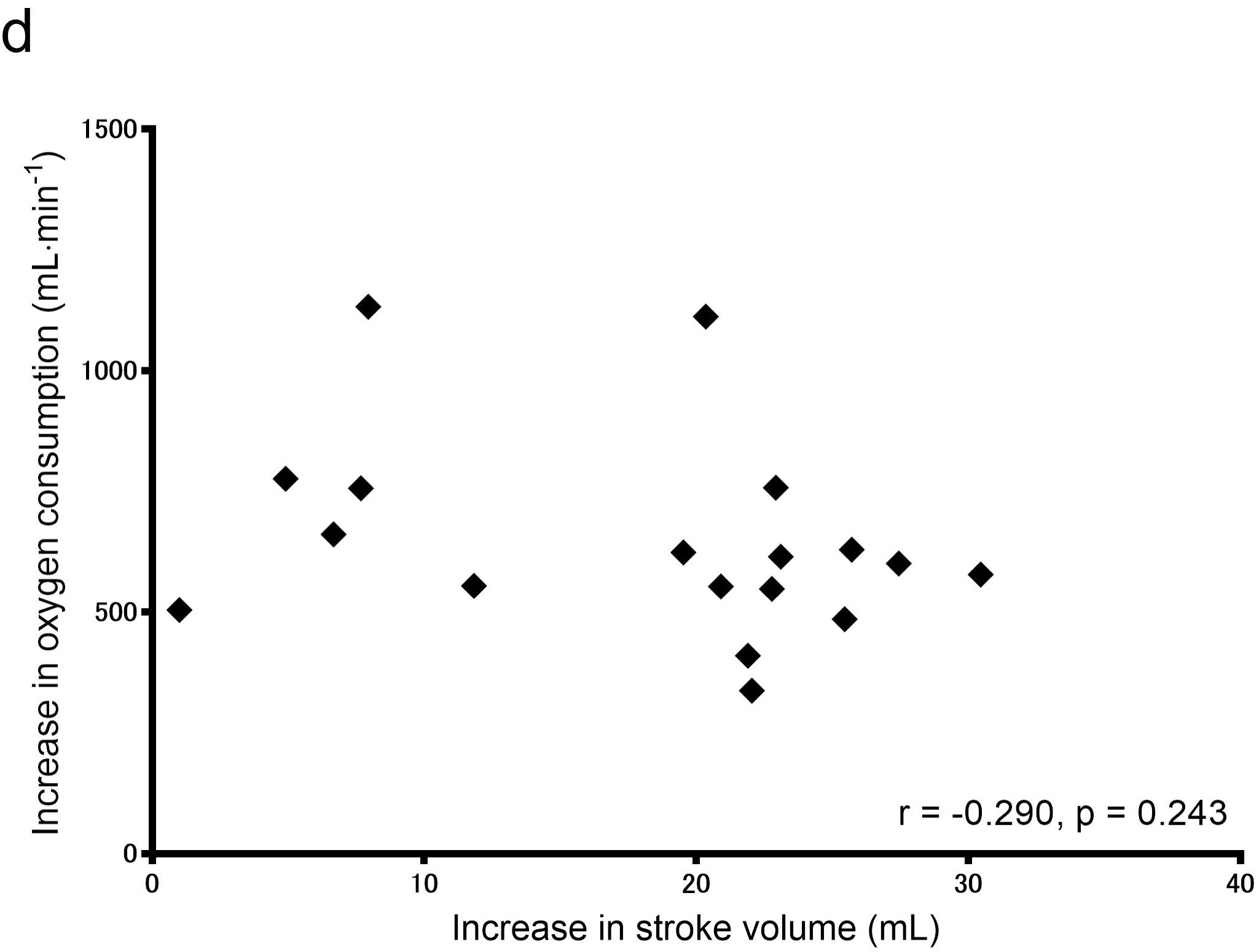
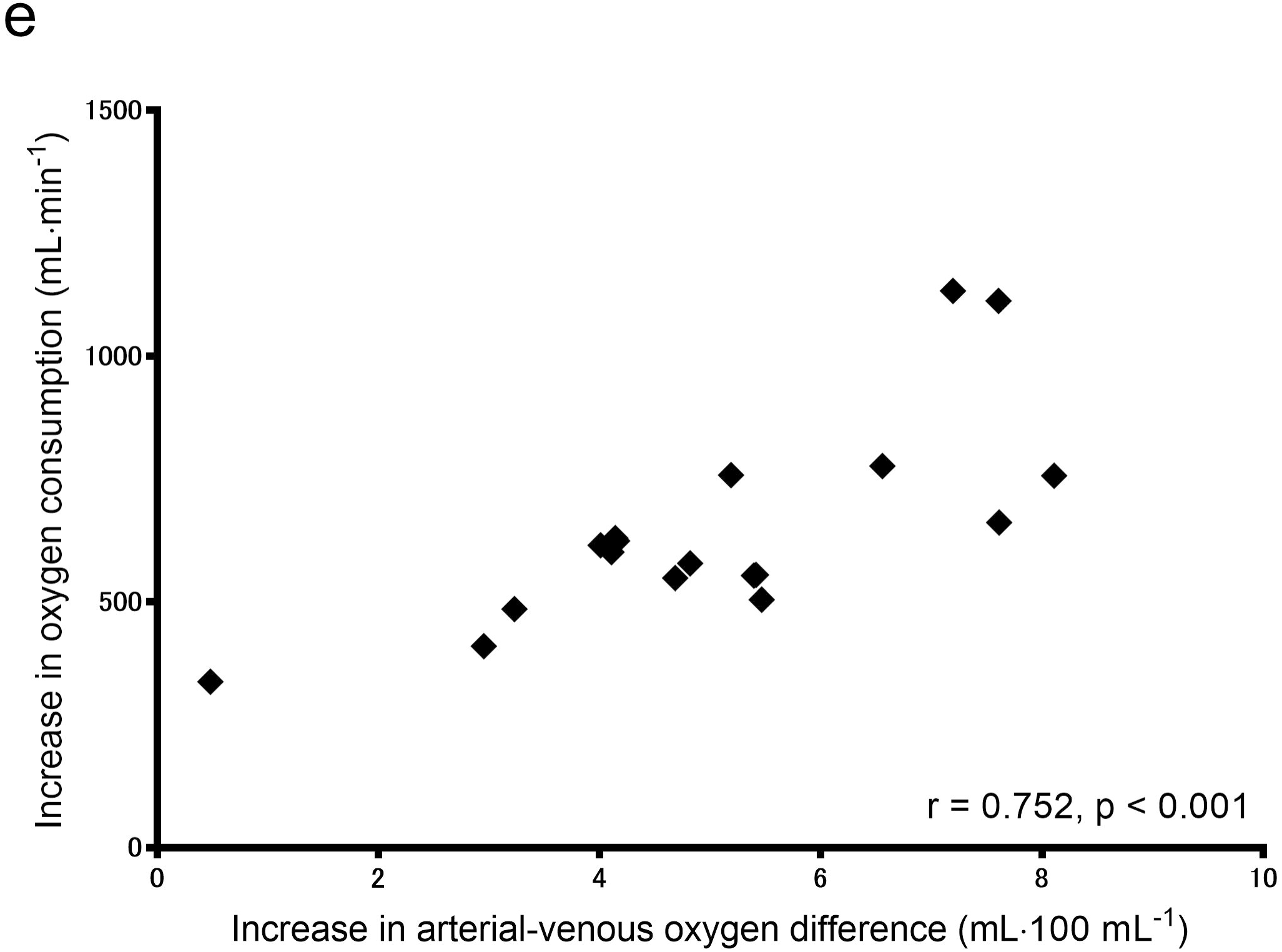
Correlations between the increases in V̇O_2_ and respiratory rate (a), V̇O_2_ and tidal volume (b), V̇O_2_ and heart rate (c), V̇O_2_ and stroke volume (d), and V̇O_2_ and arterial-venous oxygen difference (e) from rest to ventilatory threshold.

**Table 3.** Stepwise multiple regression analysis for identifying the determinants of the increase in V̇O_2_ from rest to ventilatory threshold

| Variable | $\beta$ | Coefficient | SE | t value | p value |
| --- | --- | --- | --- | --- | --- |
| Increase in heart rate from rest to ventilatory threshold | 0.546 | 8.73 | 2.78 | 3.14 | 0.007 |
| Increase in arterial-venous oxygen difference from rest to ventilatory threshold | 0.398 | 43.03 | 18.82 | 2.29 | 0.037 |
| Constant |  | 133.20 | 83.61 | 1.59 | 0.132 |
| F (2, 15) = 21.12, p < 0.001, $R^2 = 0.738$, Adjusted $R^2 = 0.703$ | | | | | |

242 $\beta$ , standard coefficient; SE, standard error.

From rest to peak exercise, correlations between the increases in V̇O_2_ and other physiological variables are shown in Fig 3. The increase in V̇O_2_ was significantly correlated with the increases in tidal volume (r = 0.806; 95% CI = 0.544, 0.925; and p < 0.001) (Fig 3b), heart rate (r = 0.691; 95% CI = 0.330, 0.875; and p = 0.002) (Fig 3c), and arterial-venous oxygen difference (r = 0.729; 95% CI = 0.398, 0.892; and p < 0.001) (Fig 3e). Stepwise multiple regression analysis revealed that the increases in tidal volume (β = 0.611) and heart rate (β = 0.353) were significant determinants for the increase in V̇O_2_ (adjusted R^2^ = 0.702, p < 0.001) (Table 4).

**Fig 3.**
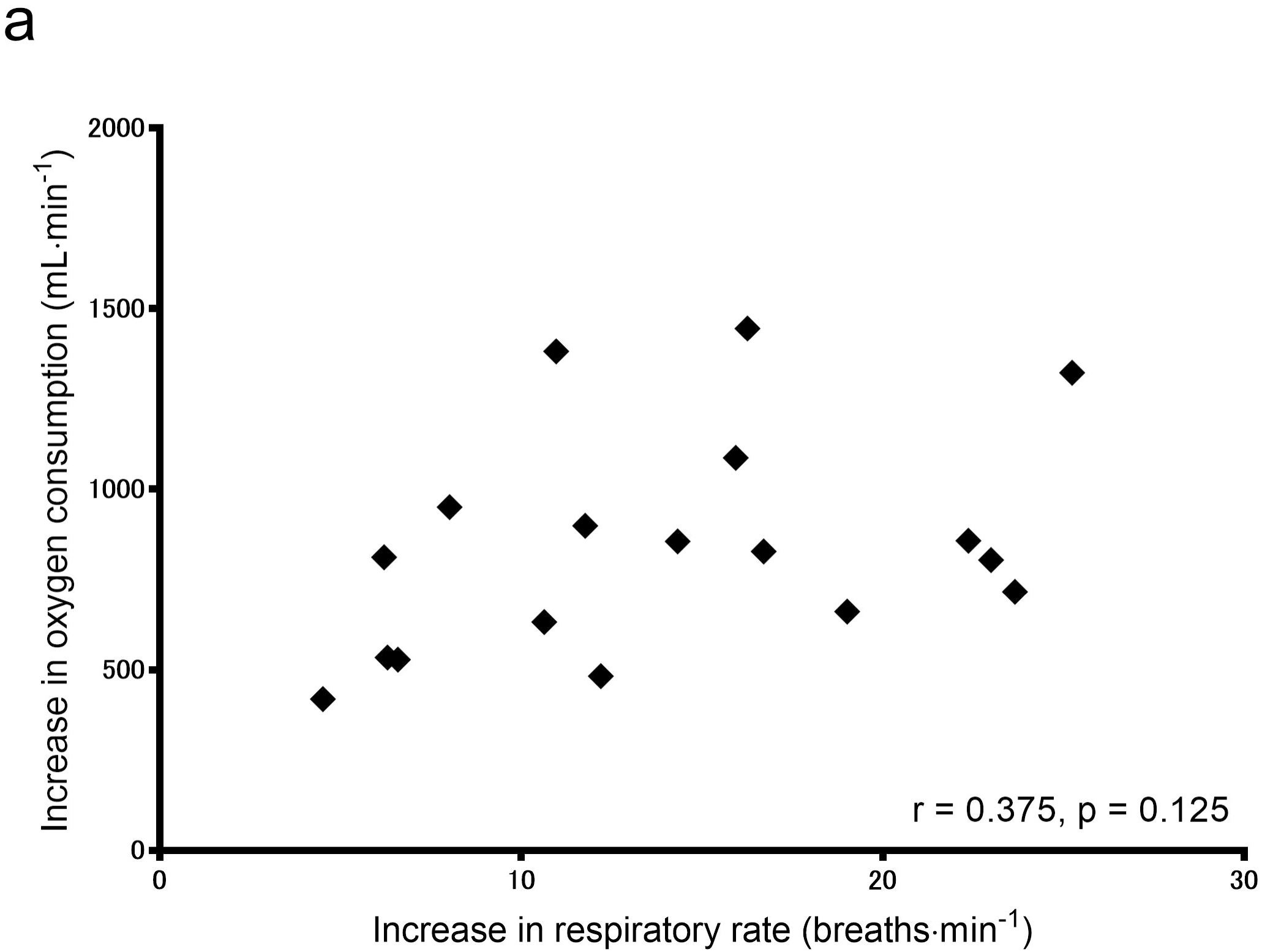
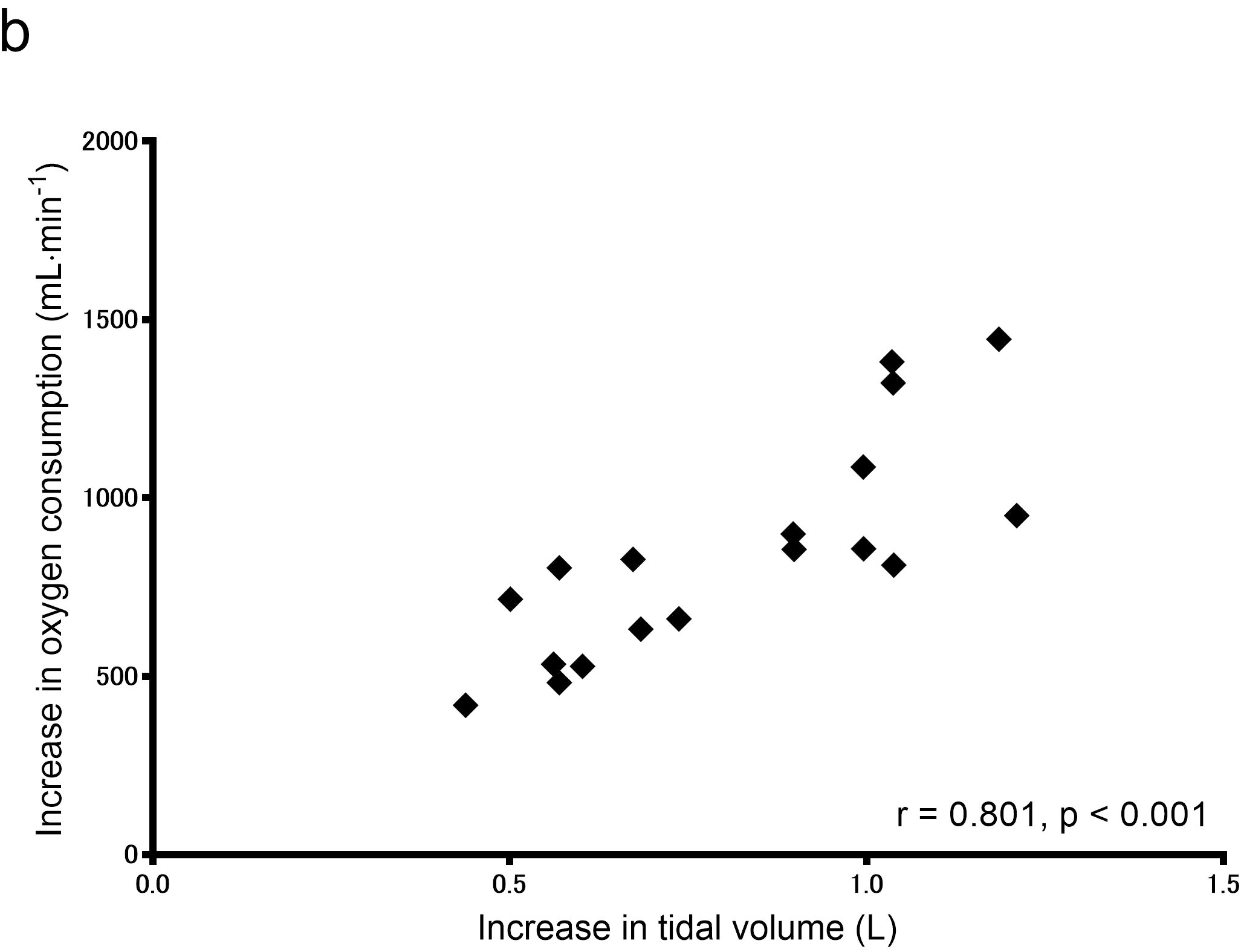
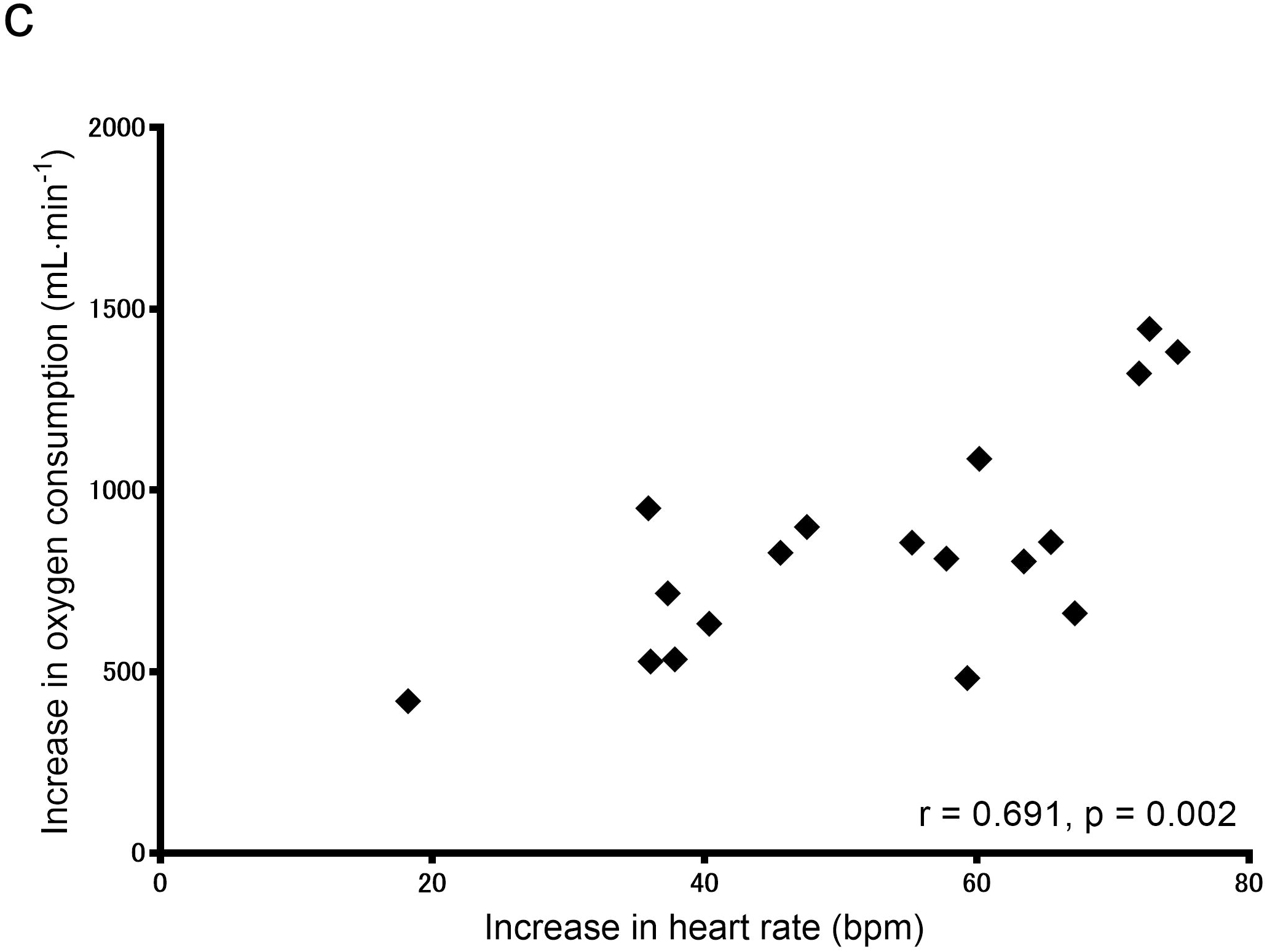
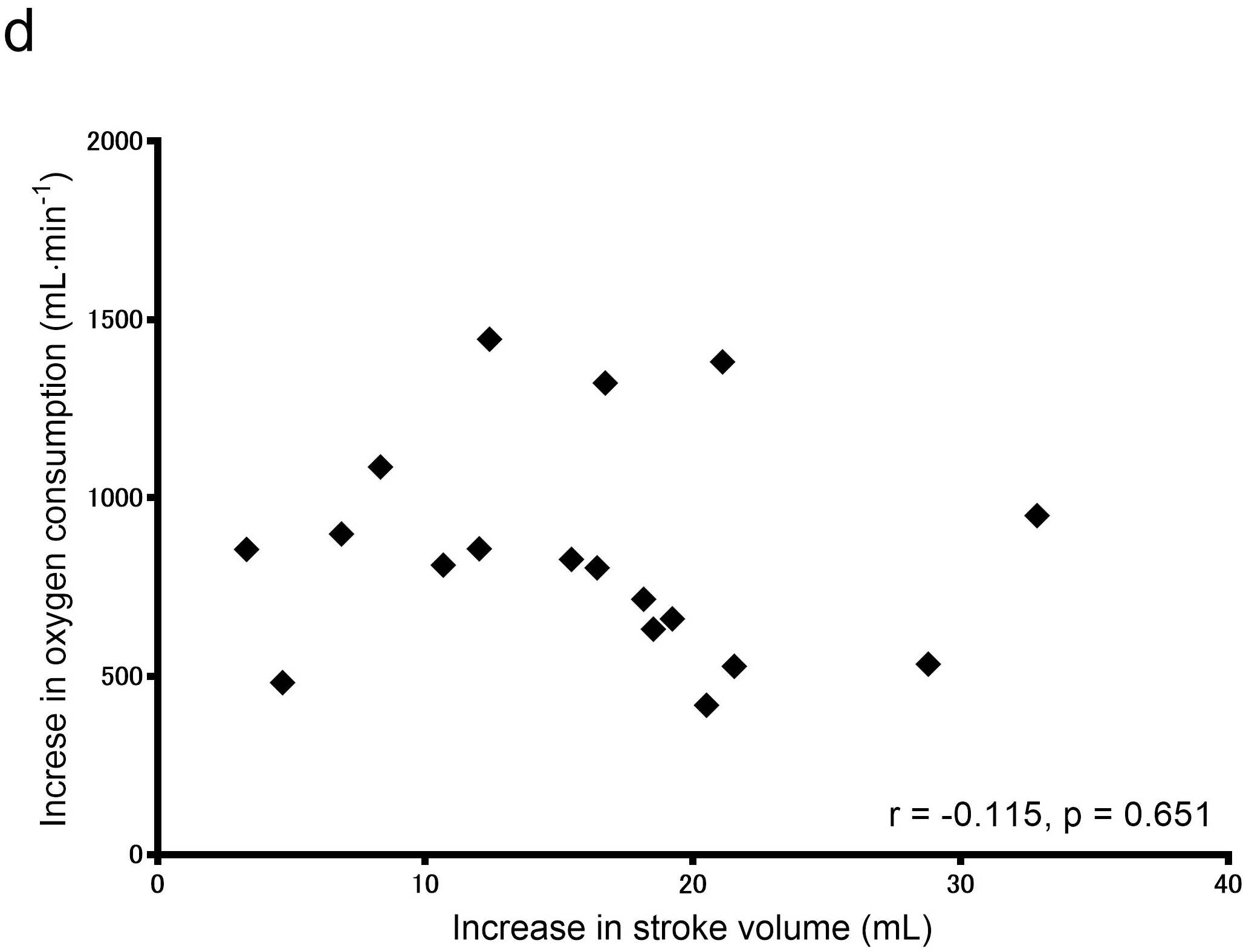
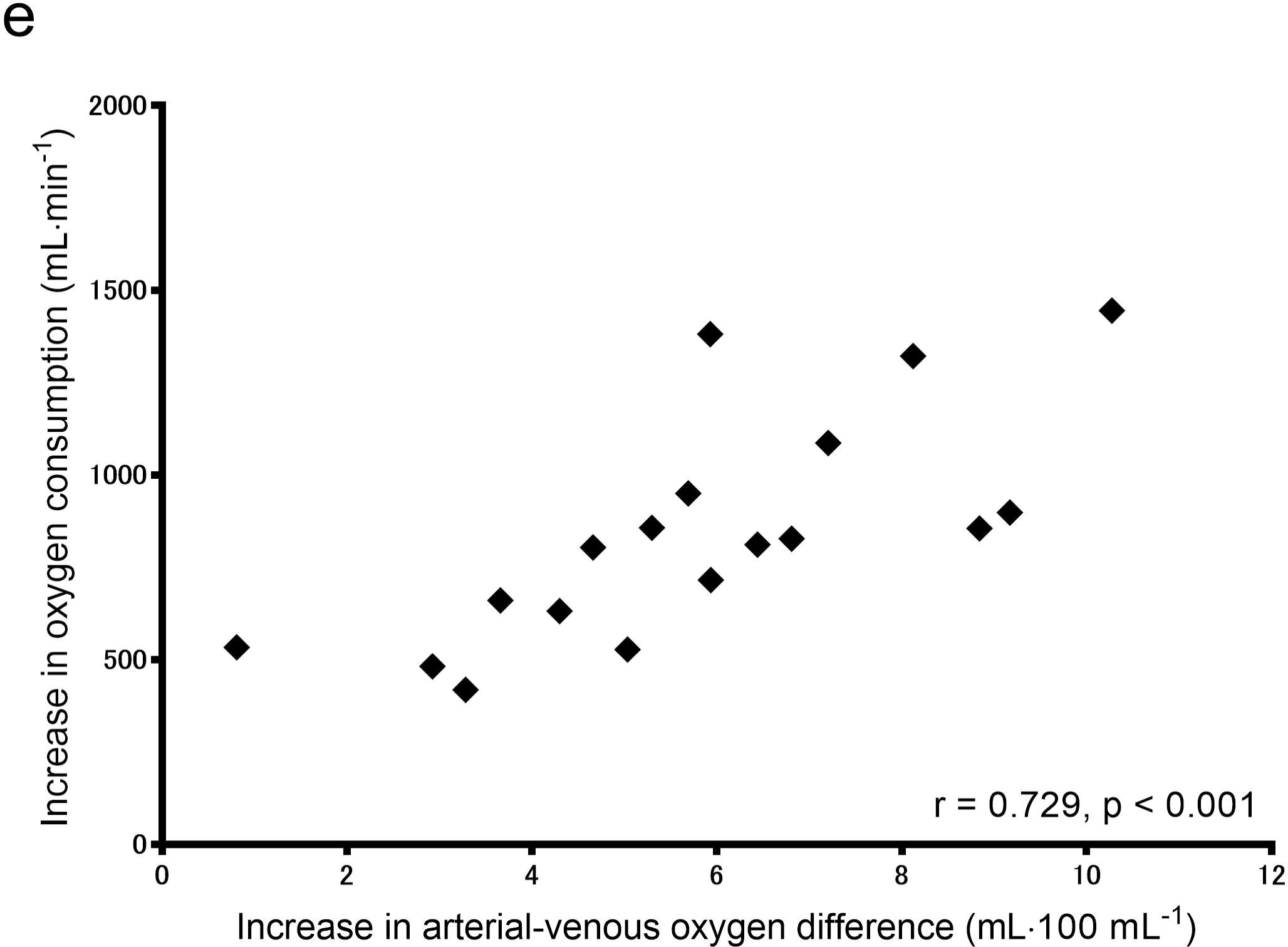
Correlations between the increases in V̇O_2_ and respiratory rate (a), V̇O_2_ and tidal volume (b), V̇O_2_ and heart rate (c), V̇O_2_ and stroke volume (d), and V̇O_2_ and arterial-venous oxygen difference (e) from rest to peak exercise.

**Table 4.** Stepwise multiple regression analysis for identifying the determinants of the increase in V̇O_2_ from rest to peak exercise

| Variable | $\beta$ | Coefficient | SE | t value | p value |
| --- | --- | --- | --- | --- | --- |
| Increase in tidal volume from rest to peak exercise | 0.611 | 749.82 | 194.88 | 3.85 | 0.002 |
| Increase in heart rate from rest to peak exercise | 0.353 | 6.72 | 3.02 | 2.23 | 0.041 |
| Constant |  | -117.65 | 155.12 | 0.76 | 0.460 |
| F (2, 15) = 20.99, $p < 0.001$, $R^2 = 0.737$, Adjusted $R^2 = 0.702$ | | | | | |

$\beta$ , standard coefficient: SE, standard error.

## Discussion

This study is the first to explore the physiological determinants of the increase in V̇O_2_ during graded exercise in individuals with stroke. From rest to ventilatory threshold, the increases in heart rate and arterial-venous oxygen difference were the significant determinants of the increase in V̇O_2_. From rest to peak exercise, the increases in tidal volume and heart rate were the significant determinants of the increase in V̇O_2_. Physiological impairments in these variables may negatively affect cardiorespiratory fitness in individuals with stroke.

The results of correlation analyses may support previous studies suggesting that impaired tidal volume, heart rate, and arterial-venous oxygen difference at peak exercise limit cardiorespiratory fitness in individuals with stroke [12, 16-18, 31]. Contrary to our hypothesis, only the increase in heart rate was identified as one of the significant determinants of both the increases in V̇O_2_ from rest to ventilatory threshold and that from rest to peak exercise, which can be explained by the fact that V̇O_2_ increases nearly linearly with increasing heart rate during exercise [35]. Tomczak et al. [18] indicated that the impaired increase in V̇O_2_ during exercise in individuals with stroke may be attributed to the impaired increase in heart rate. Our results suggest that cardiac function to supply oxygen rather than the ability of skeletal muscles to extract oxygen affects cardiorespiratory fitness in individuals with stroke. Conversely, Jakovljevic et al. [17] and Moore et al. [19] reported that oxygen extraction rather than oxygen supply is related with cardiorespiratory fitness in individuals with chronic stroke. The discrepancy in results between the present and previous studies [17, 19] may be caused by the characteristics of participants such as time since stroke.

Skeletal muscle changes after stroke, such as muscle atrophy and shift of muscle fiber type (from type I slow-twitch muscle fibers to type II fast-twitch muscle fibers) particularly in the paretic lower extremity, are observed in individuals with stroke [36]. In addition, impaired vasodilatory function and reduction in blood flow in the paretic lower extremity have also been reported [37, 38]. These changes after stroke may reduce the ability of skeletal muscles to extract oxygen. Reduction in the ability of skeletal muscles to extract oxygen during exercise may increase the dependence of anaerobic glycolysis for energy output, thus increasing the output of lactic acid [39, 40]. Minute ventilation begins to increase exponentially relative to the increase in V̇O_2_ to eliminate the excess carbon dioxide produced from bicarbonate buffering of lactic acid, which is defined as ventilatory threshold [41]. These findings support our results that arterial-venous oxygen difference was identified as one of the physiological determinants of the increase in V̇O_2_ from rest to ventilatory threshold.

Although arterial-venous oxygen difference increases with incremental exercise, that is assumed to be relatively constant at a submaximal work rate [35]. We observed a significant increase in V̇O_2_ but not in arterial-venous oxygen difference from ventilatory threshold to peak exercise. This may explain the reason why the increase in arterial-venous oxygen difference was not selected as the significant physiological determinant of the increase in V̇O_2_ from rest to peak exercise. Our stepwise regression analysis indicated that the increase in tidal volume was a major physiological determinant of the increase in V̇O_2_ from rest to peak exercise. Sisante et al. [16] and Tomczak et al. [18] reported that tidal volume and V̇O_2_ at peak exercise are significantly lower in individuals with stroke than in healthy controls. Their findings suggest that decreased tidal volume limits the increase in V̇O_2_ during exercise. The paralysis of the expiratory muscles on the affected side, decreased motion of the diaphragm, and reduced chest wall excursion may limit the increase in tidal volume during exercise [12, 42].

This study had several limitations. First, all participants were in the subacute stages of recovery from stroke. Therefore, generalization of the findings to individuals with chronic stroke should be made with caution. Second, we used a recumbent cycle ergometer. A treadmill [6], a total-body recumbent steppe [43], a robotics-assisted tilt table [30], and an arm crank ergometer [31] are also used to assess cardiorespiratory fitness in individuals with stroke. Further studies are warranted to examine whether the major physiological determinant of the increase in V̇O_2_ during exercise differs with the exercise devices. Finally, as this study used the cross-sectional observational design, the physiological determinants of the temporal changes in V̇O_2_ at ventilatory threshold and at peak exercise could not be examined. Further longitudinal studies are needed to examine whether physiological impairments in tidal volume, heart rate, and arterial-venous oxygen difference affect the temporal changes in V̇O_2_ for the development of appropriate therapies to improve cardiorespiratory fitness in individuals with stroke.

## Conclusions

V̇O_2_ is well-known to increase nearly linearly with increasing heart rate [35, 41]; however our results suggest that arterial-venous oxygen difference and tidal volume are also significant physiological determinants of increase in V̇O_2_ from rest to ventilatory threshold and that from rest to peak exercise, respectively. Our findings could potentially contribute to the development of appropriate therapies to improve cardiorespiratory fitness in individuals with stroke.

## Acknowledgments

The authors certify that no other persons have made substantial contributions to this manuscript.

